# Bacterial quorum sensing controls carbon metabolism to optimize growth in changing environmental conditions

**DOI:** 10.1101/2024.01.21.576522

**Authors:** Chelsea A. Simpson, Zach Celentano, James B. McKinlay, Carey D. Nadell, Julia C. van Kessel

## Abstract

Bacteria sense population density via the cell-cell communication system called quorum sensing (QS). Some QS-regulated phenotypes (*e.g.*, secreted enzymes, chelators), are public goods exploitable by cells that stop producing them. We uncovered a phenomenon in which *Vibrio* cells optimize expression of the methionine and tetrahydrofolate (THF) synthesis genes via QS. Strains that are genetically ‘locked’ at high cell density grow slowly in minimal glucose media and suppressor mutants accumulate via inactivating-mutations in *metF* (methylenetetrahydrofolate reductase) and *luxR* (the master QS transcriptional regulator). Methionine/THF synthesis genes are repressed at low cell density when glucose is plentiful and are de-repressed by LuxR at high cell density as glucose becomes limiting. In mixed cultures, QS mutant strains initially co-exist with wild-type, but as glucose is depleted, wild-type outcompetes the QS mutants. Thus, QS regulation of methionine/THF synthesis is a fitness benefit that links private and public goods within the QS regulon, preventing accumulation of QS-defective mutants.

## Introduction

Bacteria monitor their surroundings and respond to changes in cell density through quorum sensing (QS), allowing them to adapt to environmental cues that affect bacterial cell growth and survival^1,2^. Using molecules called autoinducers, cells sense density increases and respond with broad changes in gene expression that alter their behaviors and optimize growth, colonization, infection, and many other processes. QS signaling mechanisms have been studied in several marine *Vibrio* species^3–6^, which are a group of ecologically and economically relevant pathogens of fish, shellfish, and mammals, including humans. Early foundational QS studies were performed with *V. campbellii* strain ATCC BAA-1116 (also called BB120), a *V. campbellii* isolate that was previously classified as *Vibrio harveyi*^7^. A close relative of BB120, wild isolate *V. campbellii* DS40M4 has a similar QS circuit^8^. Cells detect and respond to autoinducer signals via multiple histidine kinases that converge to phosphorylate the response regulator LuxO (Fig. 1A). At low cell density (LCD), LuxO is phosphorylated and activates transcription of the quorum regulatory small RNAs (Qrrs) that post-transcriptionally repress translation of LuxR, the high cell density (HCD) master regulator (Fig. 1A)^9^. At high cell density (HCD), de-phosphorylated LuxO is unable to activate the Qrrs, allowing for high expression of LuxR. LuxR activates various behaviors, such as enzyme secretion, that can be utilized by other cells (public goods) (Fig. 1A).

**Figure 1:**
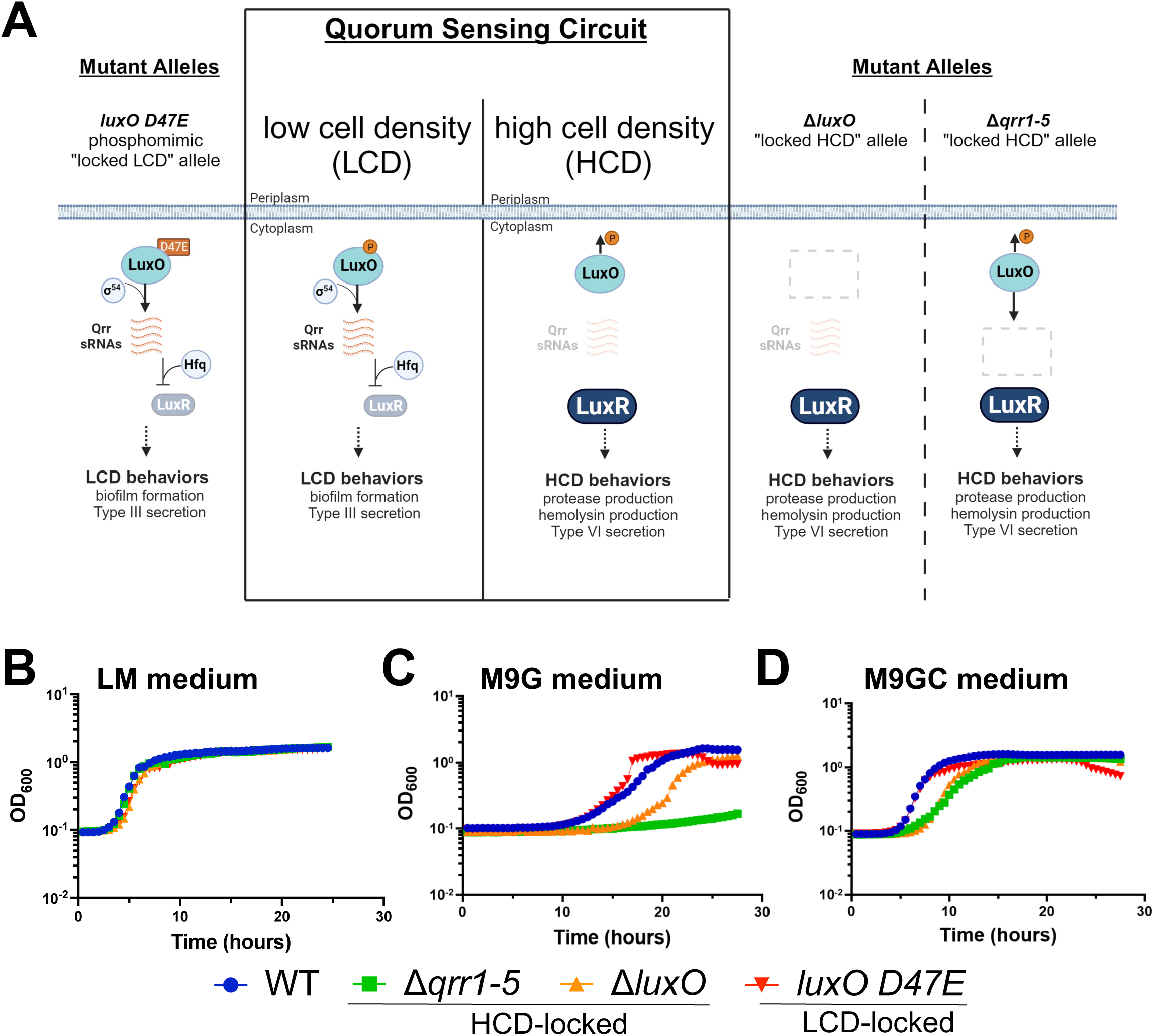
Growth trends of *V. campbellii* quorum sensing mutants differs in the absence of amino acids. (A) *V. campbellii* quorum sensing system and mutant alleles. At LCD, LuxO is phosphorylated and along with Sigma-54 activates transcription of the Qrr small RNAs. The Qrrs together with Hfq repress LuxR production. At HCD, unphosphorylated LuxO is unable to activate transcription of the Qrr small RNAs, allowing for translation of LuxR and production of proteases. The *luxO D47E* allele produces a locked-LCD phenotype. Deletion of *luxO* or the *qrrs* results in phenotypically HCD-locked strains. (B, C, D) Growth curves for *V. campbellii* DS40M4 strains in LM (B), M9G (C), or M9GC (D) media. The y-axis represents cell density OD_600_. For all panels, the data are from a single experiment that is representative of at least three independent biological experiments for each strain under every condition.

*V. campbellii* DS40M4 is completely prototrophic and grows robustly in minimal M9 medium with only the addition of carbon (*e.g.*, glucose). However, we observed that HCD-locked mutant strains of DS40M4 grew poorly in minimal media conditions compared to the wild-type parent strain, and this effect was relieved by addition of amino acids to the medium. Here, we determined the molecular mechanisms that enable QS to regulate methionine and tetrahydrofolate biosynthesis genes as cells transition between LCD and HCD states. Our results underscore the relevance of nutrient acquisition – and how it is influenced by QS regulation – in connection to the evolutionary stability of QS.

## Results

### Carbon source determines QS mutant growth trends

Because QS enables cells to respond to external signals, we sought to examine the impact of nutrients on the growth and signaling of DS40M4 strains. As previously performed in other *Vibrio* strains, we engineered mutants that are genetically locked at LCD or HCD via alleles of *luxO*: the *luxO D47E* strain constitutively activates transcription of the Qrrs and is a LCD-locked mutant, and the Δ*luxO* strain is locked at HCD and does not transcribe the Qrrs (Fig. 1A)^8^. We assayed growth of QS mutant strains in the following conditions: 1) rich, undefined <u>L</u>ysogeny Broth <u>M</u>arine medium (LM; LB with 2% NaCl), 2) <u>M9</u> minimal medium with 20 mM <u>g</u>lucose (M9G), and 3) <u>M9G</u> with <u>c</u>asamino acids (0.2%; M9GC). In this assay, the cells were washed out of LM rich medium from the overnight culture and diluted 1:100,000 into fresh medium (LM, M9C, or M9GC). Growth trends for all strains were similar in LM (Fig. 1B), and as previously observed, DS40M4 had a faster growth rate than BB120 in LM (Fig. 1B, S1A)^8^. The two HCD-locked DS40M4 strains, Δ*luxO* and Δ*qrr1-5,* did not grow well in M9G, with long lag phases that were markedly recovered by the addition of casamino acids (Fig. 1C, 1D). Trends were qualitatively similar for BB120 (Fig. S1A-C). We observed variability in the length of the lag phases for DS40M4 in the Δ*luxO* and Δ*qrr1-5* strains from experiment to experiment (Fig. S1D), suggesting that the length of the lag phase was dictated by the presence and abundance of suppressor mutants in the initial inoculum. However, in every case, the Δ*qrr1-5* strain grew the slowest in M9G (Fig. 1C, S1B, S1D), possibly because this strain completely lacks the *qrr* genes whereas the Δ*luxO* strain encodes these genes but does not activate their expression. Conversely, in both the DS40M4 and BB120 isolates, the phosphomimic *luxO D47E* strain grew robustly in M9G (Fig. 1C, S1B). We noted a consistent drop in optical density (OD) in late stationary phase in both M9G and M9GC for the *luxO D47E* strains, but that did not correlate to a decrease in cell viability (Fig. 1C, 1D, S1E, S1F). To determine if the growth defect was linked to glucose only, we assayed other sugars and observed similar growth phenotypes when DS40M4 strains were grown in either M9G or M9 with added maltose (Fig. S1G). From these data, we conclude that the presence of amino acids in the medium increases growth rates in *V. campbellii*, and in the absence of amino acids, the growth rate is dependent on the QS state.

### Deletion of metF improves growth trends of HCD strains

To identify a possible genetic link to the limited growth phenotype of the HCD-locked strains, we collected DS40M4 suppressor mutants that grew better in M9G medium than the HCD-locked parent strains Δ*qrr1-5* or Δ*luxO*. Whole genome sequencing of two suppressor mutants identified mutations in the same two loci for both suppressor strains: 1) a 305-bp deletion between genes annotated as *metL* and *metF* and a point mutation resulting in a LuxR missense allele, G37V (the Δ*qrr1-5* suppressor strain; Fig. 2A), and 2) a 25-bp deletion in *metF* and a frameshift in LuxR (the Δ*luxO* suppressor strain; Fig. S2A). To determine the individual effects of these mutations in an isogenic background strain, we introduced the following mutations in the DS40M4 Δ*qrr1-5* strain: Δ*metF*, Δ*metL*, *luxR* G37V, and/or Δ*luxR*. The Δ*qrr1-5* Δ*metL* strain did not grow in M9G medium and cells were not viable (Fig. 2B, S2F), whereas the Δ*qrr1-5* Δ*metF* strain had an increased growth rate compared to Δ*qrr1-5* (Fig. 2B, 2C). The same phenotypes occurred with the Δ*luxO* Δ*metL* and Δ*luxO* Δ*metF* strains, respectively (Fig. S2B). The *luxR* G37V allele also increased growth rates compared to the Δ*qrr1-5* parent strain (Fig. 2B, 2C) and the Δ*luxO* parent strain (Fig. S2B). The combination of both *luxR* and *metF* deletions resulted in growth rates equal to that of the Δ*qrr1-5* suppressor strain (Fig. 2C). We also observed similar growth rates with either the Δ*luxR* or *luxR* G37V allele in the Δ*qrr1-5* parent (Fig. 2C), indicating that the G37V was a loss-of-function mutation. This aligns with published studies in which substitutions in the analogous residue of the LuxR homolog, HapR, in *V. cholerae* led to decreased function^10^. Complementation of *luxR* or *metF* in their respective deletion backgrounds resulted in poorer growth, indicating that deletion of *luxR* or *metF* are indeed sufficient to increase growth in the Δ*qrr1-*5 background (Fig. S2D). Further, deletion of *luxR* or *metF* also increased growth in the wild-type background (Fig. 2B, S2E). Addition of casamino acids resulted in much faster growth rates and shorter lag phases for all strains (Fig. 2D, S2C). We note that three strains (Δ*qrr1-5*, Δ*qrr1-5* Δ*metF*, and Δ*qrr1-5* Δ*metL*) had a slower growth rate than all the other strains even in the presence of casamino acids (Fig. 2D), suggesting that HCD-locked strains expressing wild-type *luxR* may have additional growth defects under these conditions.

**Figure 2:**
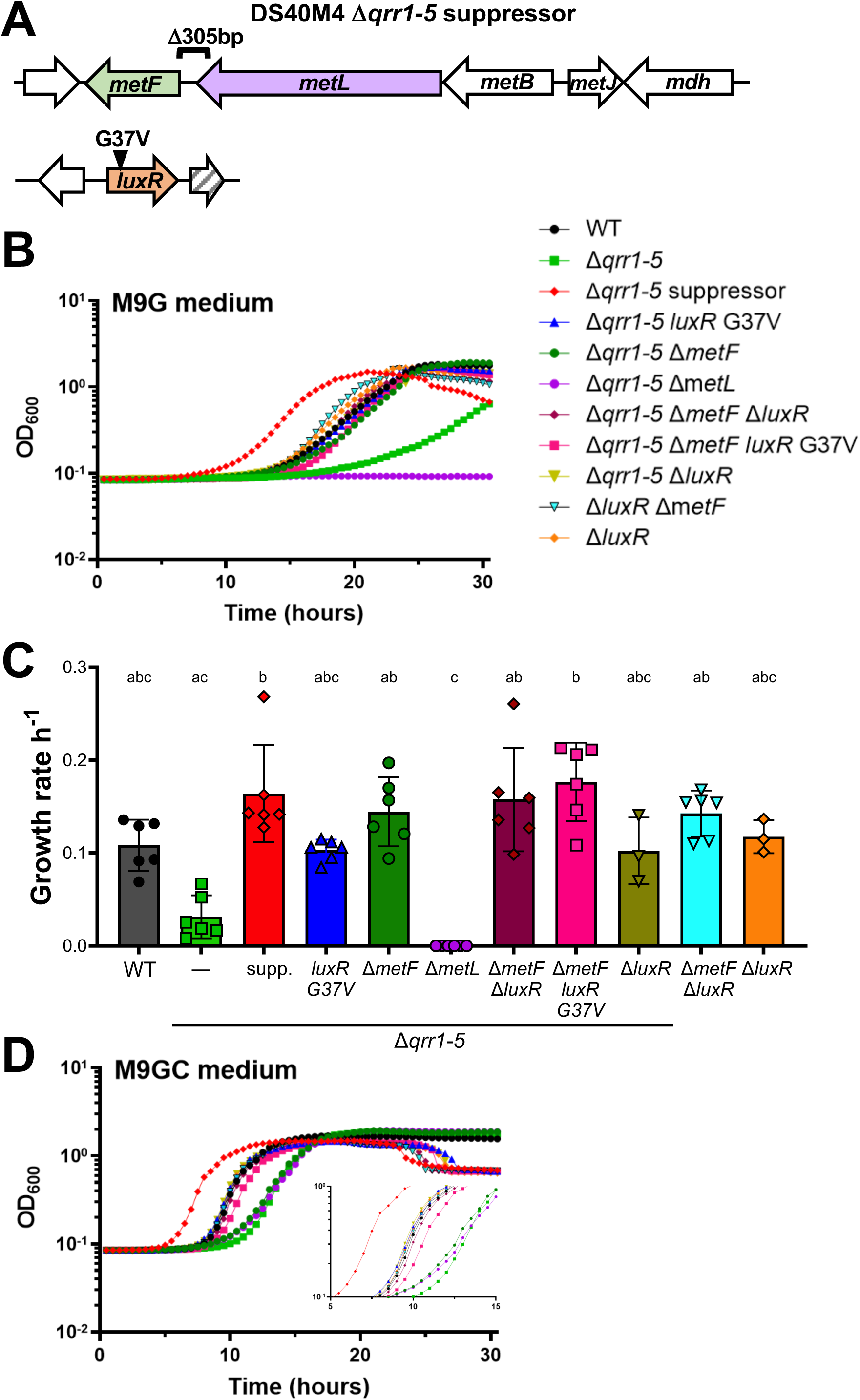
Mutation of *metF* or *luxR* restores growth to HCD-locked strains. (A) Mutations identified in the Δ*qrr1-5* suppressor mutant strain. (B) Culture growth of *V. campbellii* DS40M4 strains in M9G medium. (C) Growth rates of strains from panel B are plotted. The y-axis represents cell density OD_600._ Error bars show the mean and standard deviation of at least three biological replicates. The slope of the line was derived from the data in exponential phase with at least 5 data points per growth curve. Statistical analyses were performed using the Kruskal-Wallis test (non-parametric) followed by Dunn’s multiple comparisons test (*n =* at least *3*). Different letters indicate significance (*p<*0.05). (D) Growth curves of *V. campbellii* DS40M4 strains in M9GC medium. The y-axis represents cell density OD_600._ For panels B and D, the data are from a single experiment that is representative of at least three independent biological experiments for each strain under every condition.

### DS40M4 growth is optimal with a balance of the activated methyl cycle and folate cycle

Biochemical and genetic work in *E. coli* and other bacteria has generated a model of conserved key enzymes, substrates, and products in the methionine biosynthesis pathway, including the activated methyl cycle (AMC) and folate cycle. The KEGG^11^ database uses this information to predict the presence/absence of biosynthetic pathways present in *V. campbellii* BB120 based on gene conservation and ontology. We used this information to diagram the predicted pathways for *V. campbellii* DS40M4 (Fig. 3A). MetF (5,10-methylenetetrahydrofolate reductase) catalyzes the reduction of 5,10-methylenetetrahydrofolate (CH_2_=THF) to 5-methyltetrahydrofolate (CH_3_-THF), a step in the folate cycle that is the precursor to tetrahydrofolate (THF) synthesis (Fig. 3A). MetH and MetE are both able to use the CH_3_-THF product and L-homocysteine to produce L-methionine and THF. Methionine is then used as a substrate to produce *S-*adenosyl-L-methionine (SAM), *S-*adenosyl-L-homocysteine (SAH), and *S*-(5-deoxy-D-ribos-5-yl)-L-homocysteine (SRH)^12^. SRH is used by LuxS to regenerate L-homocysteine and the byproduct 4,5-dihydroxy-2,3-pentanedione (DPD), the precursor of autoinducer AI-2^13^.

**Figure 3:**
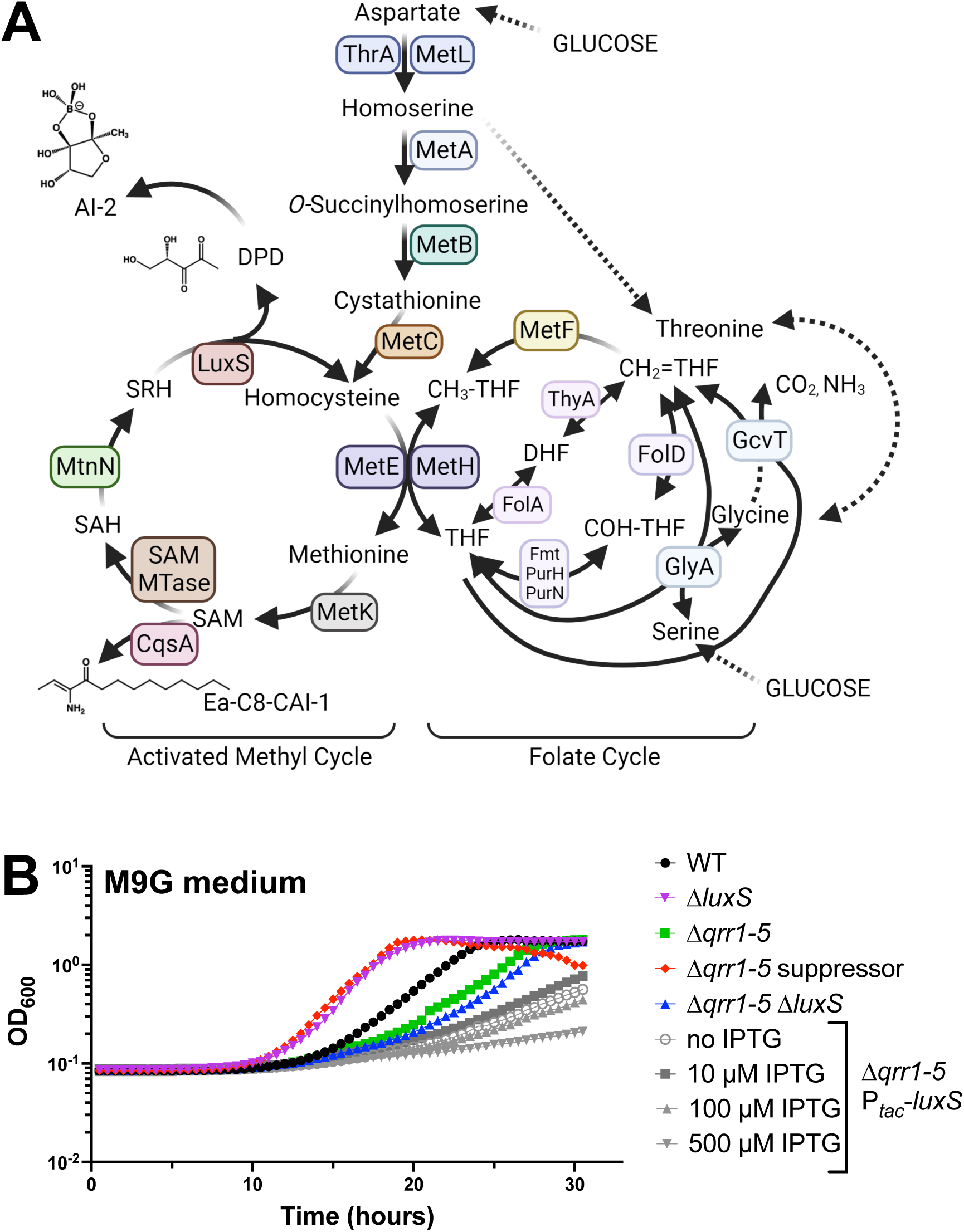
Altering flux between the activated methyl cycle and folate cycle affects culture growth. (A) Predicted activated methyl cycle and folate cycle enzymes in *V. campbellii* DS40M4 based on *V. campbellii* BB120 KEGG gene predictions. (B) Growth curves for *V. campbellii* DS40M4 strains grown in M9G medium. The y-axis represents cell density OD_600._ The data shown are from a single experiment that is representative of at least three independent biological experiments for each strain under every condition.

Based on our observation that poor growth of HCD-locked strains was alleviated by deletion of *metF*, we hypothesized that the HCD-locked strains produced higher levels of proteins involved in the AMC and/or the folate cycle. Increased MetF enzyme concentrations predictably would decrease the pool of CH_2_=THF and increase CH_3_-THF pools, possibly affecting the AMC. Thus, we next tested the effect of altering AMC substrate pools via deletion of *luxS*. The Δ*luxS* strain had increased growth in M9G medium, like the suppressor mutant (Fig. 3B). Conversely, deletion of *luxS* in the Δ*qrr1-5* background had only slightly decreased growth compared to the parent and eventually reached a similar density to wild-type (Fig. 3B, S2F). Over-expression of *luxS* via an IPTG-inducible promoter in the Δ*qrr1-5* background led to lower growth rates; higher IPTG levels correlated with decreased growth trends (Fig. 3B). Collectively, these data show that deletion of the *luxS* or *metF* gene increased growth rates, and alteration of these levels in QS-locked strains exacerbated the growth defect.

### Quorum sensing regulates transcription of methionine biosynthesis enzyme genes

In *E. coli* and other bacteria, methionine biosynthesis genes are tightly regulated: MetR, activated by homocysteine^14^, activates expression of methionine synthesis genes, and MetJ, which itself is activated by SAM^15^, opposes MetR regulation^16^ (Fig. 4A). Based on our observations that deletion of *metF* or *luxS* increased the growth rate of DS40M4, especially in HCD-locked strains, we hypothesized that HCD-locked strains grow poorly in minimal media because these strains are over-expressing the AMC and/or folate cycle genes at higher levels than wild-type. To test this hypothesis, we examined MetJ regulation of *metF*. We isolated RNA from cells grown in M9GC because not all strains grew in M9G, thus we could not compare each genotype. We observed that *metF* expression was strongly repressed by MetJ, as predicted based on the *E. coli* literature^17^ (Fig. 4B). Further, *metF* expression was significantly activated in the Δ*qrr1-5* strain compared to wild-type, which was dependent on the presence of *luxR* (Fig. 4B). Deletion of *metJ* was epistatic to all QS genotypes. Expression of *metR* mirrored that of *metF* (Fig. S3A). We had shown previously (through *in vivo* ChIP-seq and *in vitro* biochemical assays) that LuxR binds to the promoter of *metJ* in *V. campbellii* BB120 to repress its expression^18^. We confirmed that LuxR also binds the *metJ* promoter in strain DS40M4 (Fig. S3B). We conclude that LuxR directly represses *metJ*, and MetJ represses the methionine biosynthesis genes, resulting in repression of the methionine genes at LCD and de-repression at HCD.

**Figure 4:**
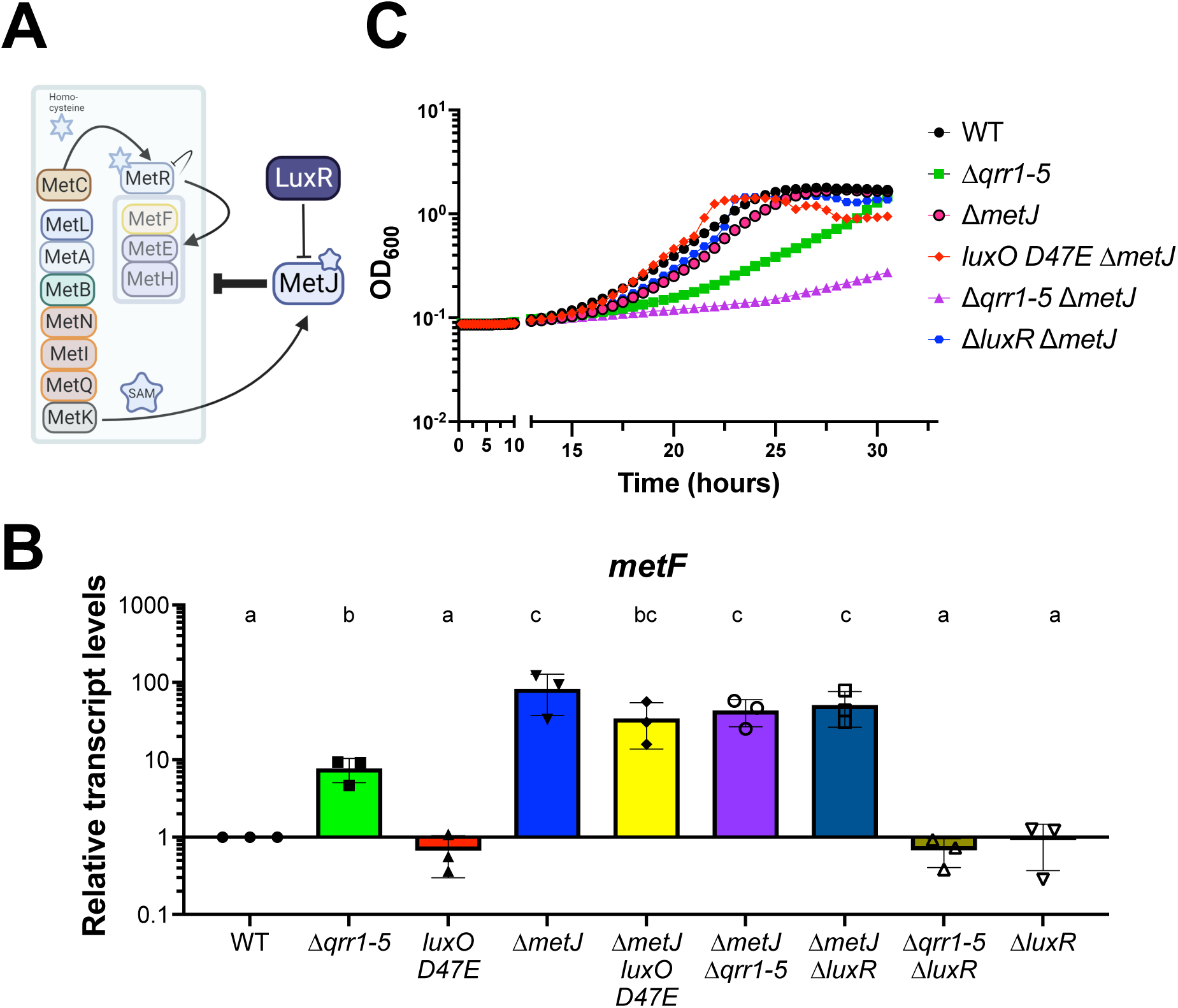
LuxR regulation of methionine biosynthesis genes. (A) Predicted regulation hierarchy of the methionine biosynthesis genes. (B) Relative *metF* transcript levels determined by RT-qPCR for cultures grown in M9GC medium. Different letters indicate significant differences *(p*<0.05). Error bars show the mean and standard deviation of three biological replicates. One-way analysis of variance (ANOVA) was performed on log-normalized data (normally distributed, Shapiro-Wilk test; *n*=3 biological replicates, Tukey’s multiple comparisons test). (C) Growth curves for *V. campbellii* DS40M4 strains in M9G medium. The y-axis represents cell density OD_600._ The data shown are from a single experiment that is representative of at least three independent biological experiments for each strain under every condition.

We hypothesized that the HCD-locked strains Δ*qrr1-5* and Δ*luxO* constitutively express the methionine biosynthesis genes even at LCD, which leads to poor growth. Thus, we next assayed the effect of deleting the MetJ master regulator on growth in minimal media. In the Δ*qrr1-5* background, deletion of *metJ* led to even more severe growth defects when grown in M9G (Fig. 4C). We observed that deletion of *metJ* does not affect growth in strains expressing the Qrrs at LCD (*e.g.,* wild-type background) (Fig. 4C). We postulated that the Qrrs might also control methionine biosynthesis genes at LCD independently of LuxR through post-transcriptional regulation. To test our hypothesis, we performed growth assays with strains expressing Qrr4 under control of an inducible P*_tac_* promoter in strain backgrounds with or without *luxR*. In M9G medium, increased Qrr4 expression improved growth in both the Δ*qrr1-5* strain and in the Δ*qrr1-5* Δ*luxR* strain (Fig. S3C). These results indicate that the Qrrs positively influence growth in glucose minimal media independently of LuxR and supports our model that the Qrrs post-transcriptionally control methionine biosynthesis genes at LCD.

### Wild-type DS40M4 strains outcompete LCD- and HCD-locked mutant strains in limited nutrients

Collectively, our observations demonstrated that *V. campbellii* DS40M4 strains that are incapable of sensing population density (*i.e.*, LCD-locked or HCD-locked strains), do not regulate methionine biosynthesis or the folate cycle optimally, leading to altered growth trends in M9G (limited nutrient conditions). We hypothesized that QS regulatory control of two key pathways – AMC and folate cycle – evolved to optimize growth in poor nutrient conditions. Thus, we assayed the fitness of wild-type DS40M4 compared to our genetically engineered *luxO D47E* (LCD-locked) and the Δ*qrr1-5* (HCD-locked) strains. First, we competed wild-type and the *luxO* D47E strain in M9G batch cultures from a range of initial population frequencies. The wild-type outcompeted the *luxO D47E* strain at every ratio tested (Fig. 5A).

**Figure 5:**
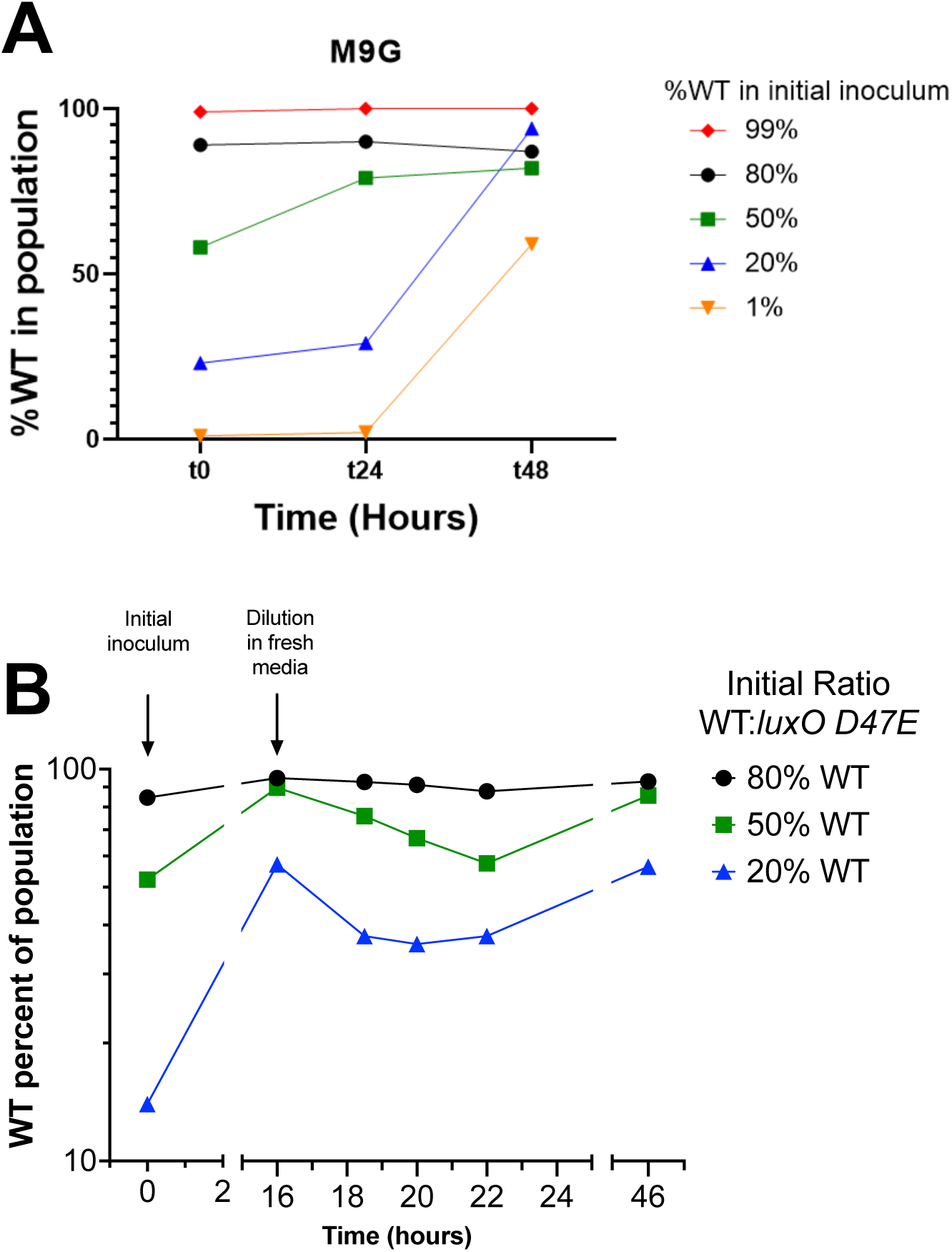
Wild-type cells outcompete cheater strains in co-culture in minimal medium. (A) Wild-type (marked with Tm) and *luxO D47E* (marked with Spec) strains of DS40M4 were inoculated at initial frequencies from 1-99% wild-type (WT) in M9G medium and allowed to grow for 48 hours. **(B**) Wild-type (marked with Tm) and *luxO D47E* (marked with Spec) strains of DS40M4 were inoculated at initial frequencies of 80%, 50%, or 20% wild-type (WT) in M9G medium and grown to near stationary phase at t=16 hours, and then diluted to maintain growth in log-phase. Colony forming units (cfu/mL) were measured at each timepoint using selective antibiotics for each strain. The total population number was calculated, and the data are represented as the percent of WT cells in the total population of WT and *luxO D47E*. The data are representative of three biological experiments.

Growth on glucose in M9G requires enzymes that are inherently not shared resources. Conversely, growth on substrates such as casein requires production of extracellular proteases to digest the casein into peptides and amino acids that are also utilizable by neighboring cells. In *Vibrio* species including *V. campbellii*, production of proteases is often positively controlled by QS^19–21^, leading to group production of amino acids that, depending on flow environment, can be shared as “public goods”. Thus, LCD-locked mutants that do not produce proteases cannot grow on their own (Fig. S4A), but they can potentially exploit the products of protease-producing cells. In stark contrast to our results competing wild-type and the *luxO* D47E strain in M9G medium, when grown in M9 with casein, the *luxO* D47E strain was able to outcompete wild-type at most ratios (Fig. S4B), indicating that wild-type production of proteases was exploited by the LCD-locked mutant, allowing it to thrive in the population. However, when there is a source of readily available amino acids (casamino acids) in the media, the wild-type and LCD-locked mutant population frequencies were maintained, suggesting a similar fitness (Fig. S4C). This short-term competition result aligns with several published *in vitro* evolution studies of the factors influencing success or failure of *V. campbellii* QS-mutants in mixed cultures^22–24^.

Because we observed that the *luxO* D47E strain could grow robustly in M9G medium but could not outcompete wild-type in that condition in a co-culture, we hypothesized that the trends are primarily due to glucose availability. Thus, we performed an experiment in which we assayed the effect of fresh nutrients on the co-cultures after initial growth. We inoculated M9G medium, allowed the co-cultured strains to grow to near stationary phase, then diluted into fresh M9G medium to keep them in log-phase growth. At each point of dilution for all three ratios, the wild-type proportion decreased (Fig. 5B). As the cells neared stationary phase again, the wild-type proportion increased for all three ratio cultures. This pattern was observed in each independent biological experiment (Fig. S4D, S4E) and suggests that 1) at LCD when there was abundant fresh carbon (glucose) present, the LCD-locked strain was able to outcompete the wild-type strain, and 2) as the cell density neared stationary phase, wild-type was able to make up for its log-phase losses and dominate the population, which we hypothesize occurred because carbon was nearing depletion. Collectively, these results indicate that the type and availability of nutrients drives the frequency of QS-defective strains in co-culture conditions.

## Discussion

The QS field has broadly studied the evolution of QS and its maintenance or loss in bacterial communities and whether cells have evolved to use different nutrient sources that might be present under differing flow conditions^25^. A key focus is the appearance and prevalence of QS null mutants – cells that do not participate in signaling, sensing and/or responding but still benefit from QS-regulated behaviors occurring at the population level^19^. Here we show that the growth rate of *V. campbellii* DS40M4 in minimal media lacking amino acids is dependent on the QS state. HCD-locked mutants exhibited poor growth in minimal media with only glucose as the carbon source, which was alleviated in suppressor mutants. Our data suggest that HCD-locked mutants grow poorly under these conditions because they express genes involved in the AMC and folate cycle at high levels, even at LCD and lag-phase. In long-term evolution studies, *V. campbellii* BB120 optimizes growth under conditions where public goods (proteases) are required for nutrient acquisition (casein); mutants that either don’t cooperate or constitutively produce public goods are less fit than wild-type in long-term serially bottle-necked experiments^22^. In addition, populations that disperse through motility, escape QS mutants and maintain cooperation^24^. In *V. campbellii* DS40M4, LCD-locked cells maintained the advantage and outcompeted wild-type when grown in minimal media supplemented with a source of fresh glucose. However, as the nutrients in the environment were depleted, wild-type cells outcompeted these LCD-locked cells. We propose this is because LCD-locked cells were incapable of QS and thus unable to activate the *met* genes to optimize glucose utilization. Conversely, in media with casein – requiring extracellular digestions with secreted proteases – LCD-locked cells were able to outcompete wild-type unless the frequency of wild-type in the population was very low. In this ratio, wild-type likely produced a limiting amount of digested casein that barely, if at all, supported growth of itself and the LCD-locked strain.

The two suppressor mutants isolated were instrumental in revealing the regulatory network connected by LuxR, the Qrrs, and the AMC and folate pathways. We propose a model that at LCD in minimal media, wild-type cells decrease, but don’t eliminate, expression of genes involved in the folate cycle and AMC and, as they transition to HCD and deplete glucose, LuxR de-represses these genes via repression of *metJ* to produce more methionine and THF (Fig. 6). Because we identified direct regulation of *metJ* via LuxR, we propose that QS regulates the flux between the AMC and folate pathways to optimize growth in nutrient limiting conditions. Such regulatory connections between QS and various metabolic processes are beginning to unfold in other systems^26,27^. In *V. cholerae*, the Qrrs alter the flux of pyruvate metabolism by repressing the translation of AlsS, which is needed to convert fermentable carbon sources into neutral products avoiding acidosis^28^. In *Burkholderia* species, methionine biosynthesis genes (*e.g.*, *metE, metF*, and *ahcY*) are activated by QS signals^29^. Thus, our study provides a new basis for studying QS regulation of glucose catabolism.

**Figure 6:**
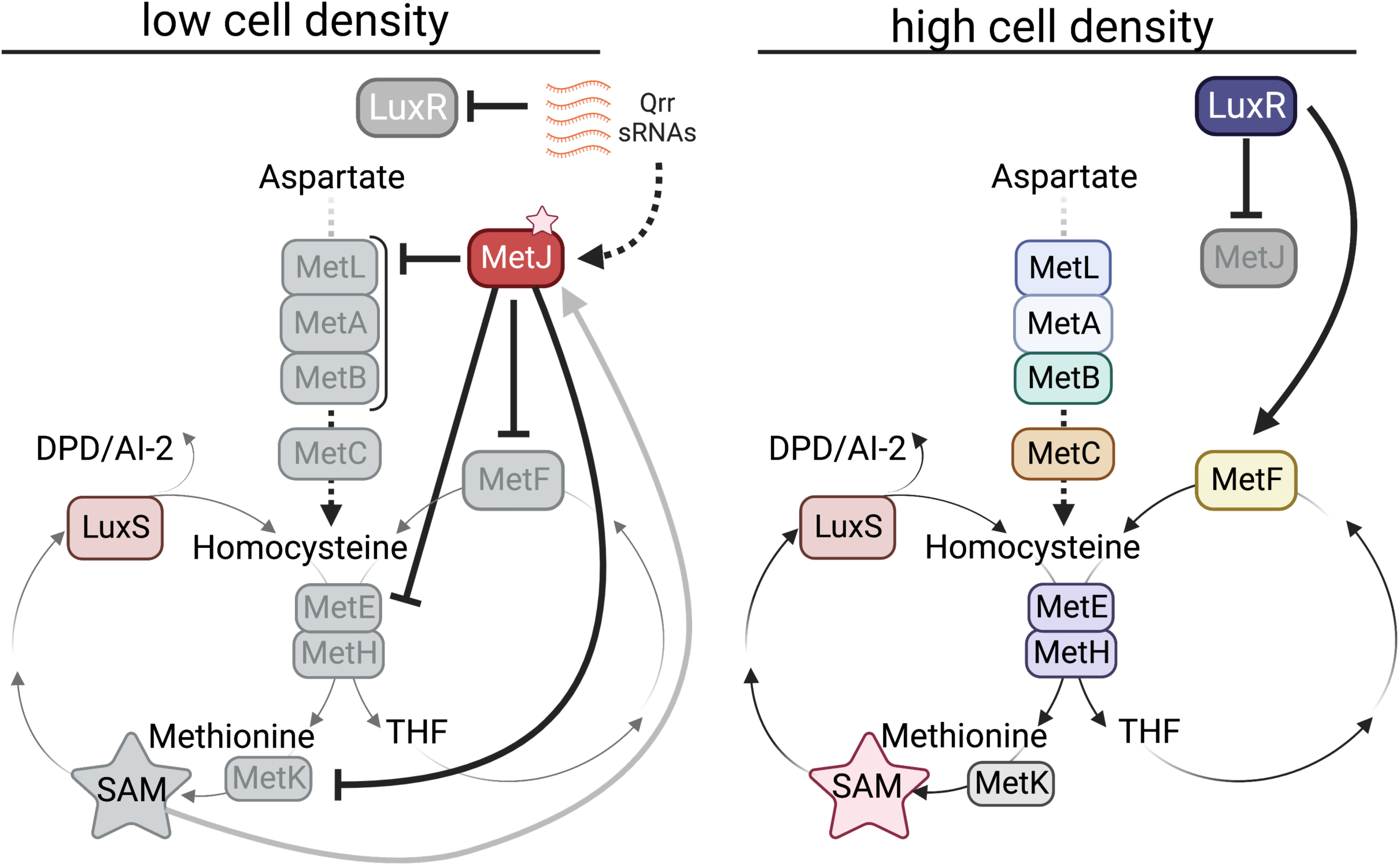
Model of quorum sensing regulation of methionine flux in M9G. At LCD, wild-type cells repress genes involved in the folate cycle and AMC and, as they transition to HCD and deplete glucose, LuxR de-represses these genes to produce methionine and THF.

Broadly, we postulate that this *V. campbellii* DS40M4 strain acquired a LuxR binding site in the *metJ* promoter that benefitted the mutant, perhaps under limited nutrient conditions, which has been maintained over time. Metabolism regulation via QS could benefit *V. campbellii* in its multiple native environments. As a planktonic free-living organism, *V. campbellii* likely has limited nutrient sources aside from complex carbohydrates (*e.g.*, chitin) available in marine ecosystems. However, during host infection (*e.g.*, shrimp, fish), *V. campbellii* likely has access to a plethora of nutrients, certainly including digestible proteins. In limited nutrient environments, we hypothesize that QS enables *V. campbellii* to optimize flux of essential biosynthesis pathways, outcompete possible competitors (QS mutants), and sustain growth. However, in nutrient-rich environments, such as the host, perhaps competition with QS mutants or other microbes may impact virulence, thus requiring cells to couple metabolism with production of public goods. Indeed, in a mouse infection model of *Vibrio cholerae*, mutant strains with deletions in either *metJ*, *metR*, or *glyA* had severe colonization defects^30^, underscoring the importance of nutrient utilization and flux regulation in different environments.

Cooperative behaviors are coordinated by QS, yet they are inherently subject to QS mutants that exploit public goods production. In several bacterial model systems, QS null mutants are isolated in wild environments; for example, in *Pseudomonas aeruginosa*, mutants lacking LasR are often isolated from infected patients^31^. However, it is yet unclear if wild populations of *V. campbellii* or other more closely related *Vibrio* species consist of QS mutants. *Vibrio* QS signals are produced by AMC enzymes and substrates, including LuxS/SRH, LuxM/SAM, and CqsA/SAM^32–34;^ methionine is converted to SAM, which is ultimately converted into autoinducers. In *V. campbellii* BB120, autoinducer concentrations increase as cell density increases^35^. However, increased production of AMC and folate cycle enzymes may also be a direct benefit as cell density increases and nutrients are depleted. Thus, we propose that *V. campbellii* is an example of a QS system that regulates private goods (methionine and THF) as both a direct benefit and because the key metabolite involved is itself inherently connected to QS signal production.

We also observed that the relative advantage or disadvantage of the *luxO D47E* mutation is highly dependent on prevailing nutrient conditions. Though QS null mutants can exploit wild-type in certain niches, QS is maintained overall in this instance because the ability to adjust to different environmental stimuli outweighs the cost of being occasionally exploited. Future studies of the genetic composition of planktonic and host-colonized *Vibrio* strains will provide important context for the evolution and maintenance of QS in *Vibrios* and other bacteria.

## Experimental Procedures

### Bacterial strains and media

*V. campbellii* strains (Table S1) were grown at 30°C on Lysogeny broth marine (LM) medium (Lysogeny broth (LB) supplemented with an additional 10 g NaCl L^-1^) or in M9 minimal media supplemented with 20 mM glucose, 0.5% casein, and 0.2% casamino acids. *E. coli* strains (Table S1) were grown in LB. Where appropriate, *Vibrio* and *E. coli* strains were supplemented with kanamycin (100 μg ml^-1^), spectinomycin (200 μg ml^-1^), gentamicin (100 μg ml^-1^) or trimethoprim (10 μg ml^-1^). Plasmids were transferred from *E. coli S17-1*λ*pir* to *Vibrio* strains by conjugation on LB plates. Exconjugants were selected on LM plates with polymyxin B at 50 U ml^-1^ and the appropriate selective antibiotic.

### Strain construction

Chitin-independent transformations were performed in DS40M4 according to Simpson et. al. 2019^37^. Strain construction details are available upon request. All genome or plasmid genotypes were confirmed by Sanger sequencing.

### Growth curves

Overnight cultures were grown in LM in culture tubes, 500 µl of cells were spun down, resuspended in 500 µl M9 medium, spun down again, and resuspended in 500 µl M9 then diluted 1:100,000 in 200 μl media. OD_600_ was measured every 30 min for 30 h using the BioTek Cytation 3 Plate Reader or BioTek Synergy H1 Plate Reader (temperature: 30°C).

### Suppressor strain isolation and sequencing

The *Vibrio* parent strain was streaked for isolation on LM plates. Colonies were grown in M9G shaking at 30°C at 275RPM until turbid. Cultures were then passaged by diluting 1:1,000 into fresh M9G until visible robust growth was seen, which was 12 passages for Δ*qrr1-5* and 17 passages for Δ*luxO*.

Sequencing and analysis of suppressor strain genomes was performed by the Indiana University Center for Genomics and Bioinformatics. Two methods were used for Breseq analysis of sequencing reads. Method 1: reads were trimmed using fastp (version 0.20.1) with parameters “-l 17 --detect_adapter_for_pe -g -p” ^39^. Breseq pipeline 0.36.^40^ with bowtie2-2.4.^41^ 2 was run with default parameters for mutation calling and annotation by re-querying against the *Vibrio campbellii* strain DS40M4 genome assembly (GenBank accession: GCA_003312585.1). Method 2: reads were adapter trimmed and quality filtered using Trimmomatic 0.38^42^ with the cutoff threshold for average base quality score set at 20 over a window of 3 bases and requiring a read length of at least 20 bp after trimming. Breseq pipeline 0.37.0^40^ with bowtie2-2.4.2 ^41^ was run with default parameters for mutation calling and annotation by re-querying against the *Vibrio campbellii* strain DS40M4 genome assembly (GenBank accession: GCA_003312585.1). The SRA accession number is SRP482837. The BioProject accession number for this data is PRJNA1063211.

### Competition assays

Strains were grown overnight in LM in culture tubes. The next day, 500 µl of cells were pelleted by centrifugation, resuspended in 500 µl M9 medium, spun down again, and resuspended in 500 µl M9, and strains were diluted 1:50 in minimal M9 media supplemented with either 20 mM glucose, 0.5% casein, or 0.2% casamino acids and grown at 30°C shaking until ∼0.5 OD_600_. Strains were adjusted to 0.5 OD_600_ with fresh media. Cultures were mixed at the desired population ratios, diluted 1:10,000, and grown at 30°C shaking at 275 RPM. Colony forming units (cfus/mL) were measured by plating serial dilutions on selective agar media. For competition experiments, where cultures were maintained in exponential phase, cultures were diluted to 0.05 OD_600_ at the 16- and 40-hour timepoints.

### RNA extraction and reverse transcriptase quantitative PCR (RT-qPCR)

RNA was extracted and qRT-PCR was performed as previously described in Simpson, et. al 2021^8^ with the exception that the strains were grown in M9GC and collected at an ∼1 OD_600_ Primers in Table S3.

### LuxR protein purification

Both WT and G37V LuxR from *V. campbellii* DS40M4 were overexpressed in *E. coli* BL21(DE3) using plasmids pZC003 and pZC004 (Table S2), respectively, derived from pET28b (Twist Bioscience). Overexpression and purification were performed as described in Newman and van Kessel. 2021^43^. The only exceptions being the use of 2.5 mL of 10x FastBreak Cell Lysis Reagent (Promega) to lyse the 1 L cell pellets and the Gel Filtration Buffer (25 mM Tris-HCl pH 8, 500 mM NaCl, and MilliQ H_2_O) used to elute the proteins. Peak eluted fractions were pooled and analyzed using SDS-PAGE. The pooled fractions were then used in P*_metJ_* EMSAs.

### EMSAs

The DNA labeling and EMSAs were performed as described in Newman and van Kessel 2021^43^. The P*_metJ_* DNA substrate was created by PCR amplifying the 239 bp segment from *V. campbellii* DS40M4 gDNA using primers ZC011 and ZC012. Reactions were run with 0.6 nM of radiolabeled DNA substrate of P*_metJ_*. The gels were dried for 20 min at 80°C and exposed to a phosphor screen for 18-22 hours. Phosphor screens were imaged on a Typhoon 9210 (Amersham Biosciences). Primers in Table S3.

## Acknowledgments

The authors thank the van Kessel lab and McKinlay lab groups for comments on the manuscript and helpful discussions. Research reported in this publication was supported by the National Institute of General Medical Sciences (NIGMS) of the National Institutes of Health (NIH) under award number R35GM124698 to JVK. The content is solely the responsibility of the authors and does not necessarily represent the official views of the National Institutes of Health. CDN is supported by the Human Frontier Science Program (award number RGY0077/2020), the Simons Foundation (award number 826672), NSF IOS grant 2017879, and NIH NIGMS grant 1R35GM151158-01. JBM is supported by the National Science Foundation CAREER award MCB-1749489.

