## Supplemental material for "Bacterial quorum sensing controls carbon metabolism to optimize growth in changing environmental conditions"

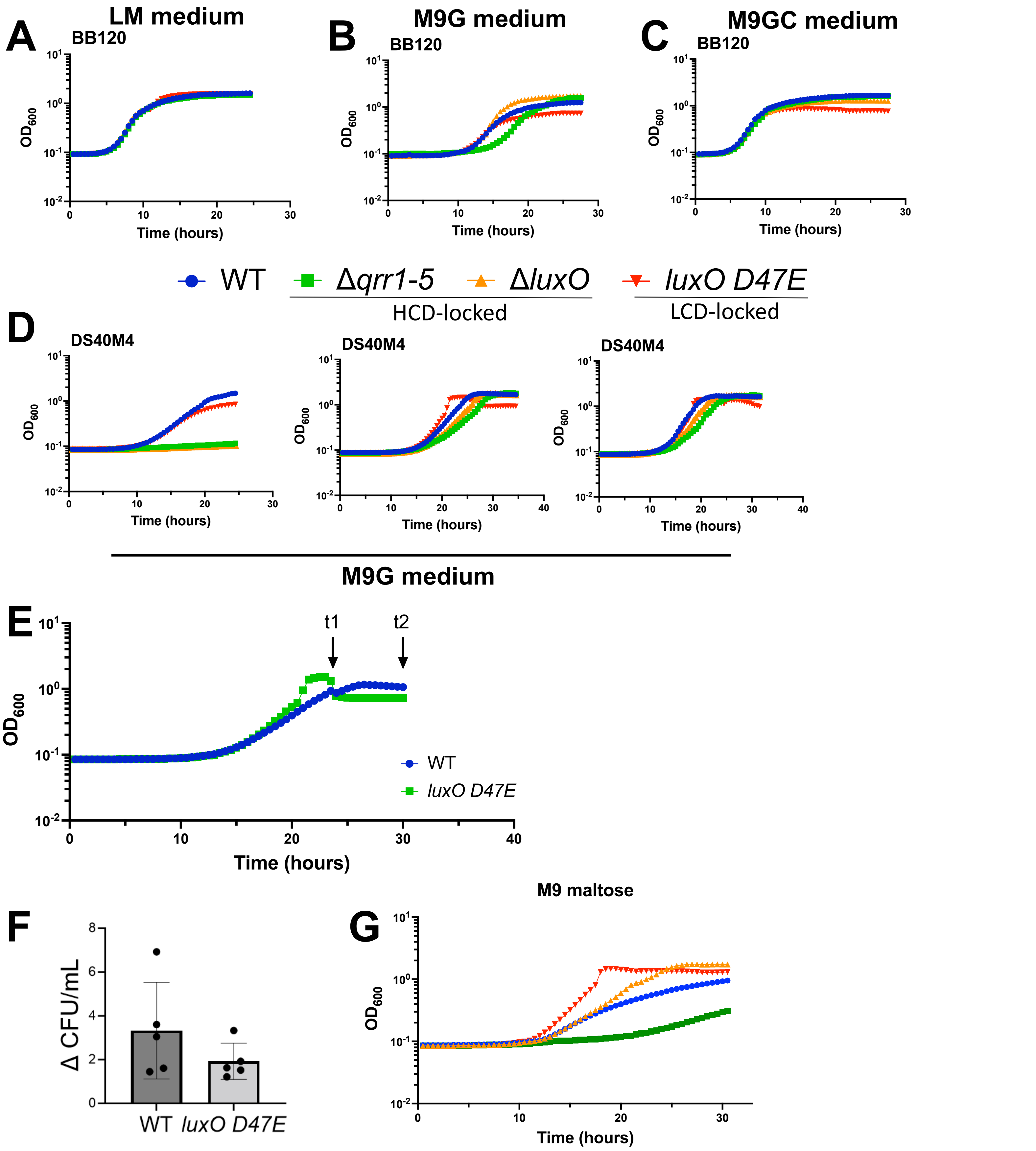

**Figure S1:** (A, B, C) Growth curves for *V. campbellii* BB120 strains in LM (A), M9G (B), or M9GC (C) media. For all panels, the data are from a single experiment that is representative of at least three independent biological experiments for each strain under every condition. (D) Variability in lag phase of HCD-locked DS40M4 mutants grown in M9G medium. Three additional biological replicates of experiments in Figure 1C are shown. (E, F) The difference in cfus/ml between timepoints 1 and 2 (E) of wild-type and *luxO D47E* DS40M4 strains grown in M9G (F). Error bars show the mean and standard deviation of five biological replicates. (G) Growth curves for *V. campbellii* DS40M4 strains in M9 medium supplemented with maltose (10 mM). (A-E,G) The y-axis represents cell density $OD_{600.}$ The data shown are from a single experiment that is representative of three independent biological experiments.

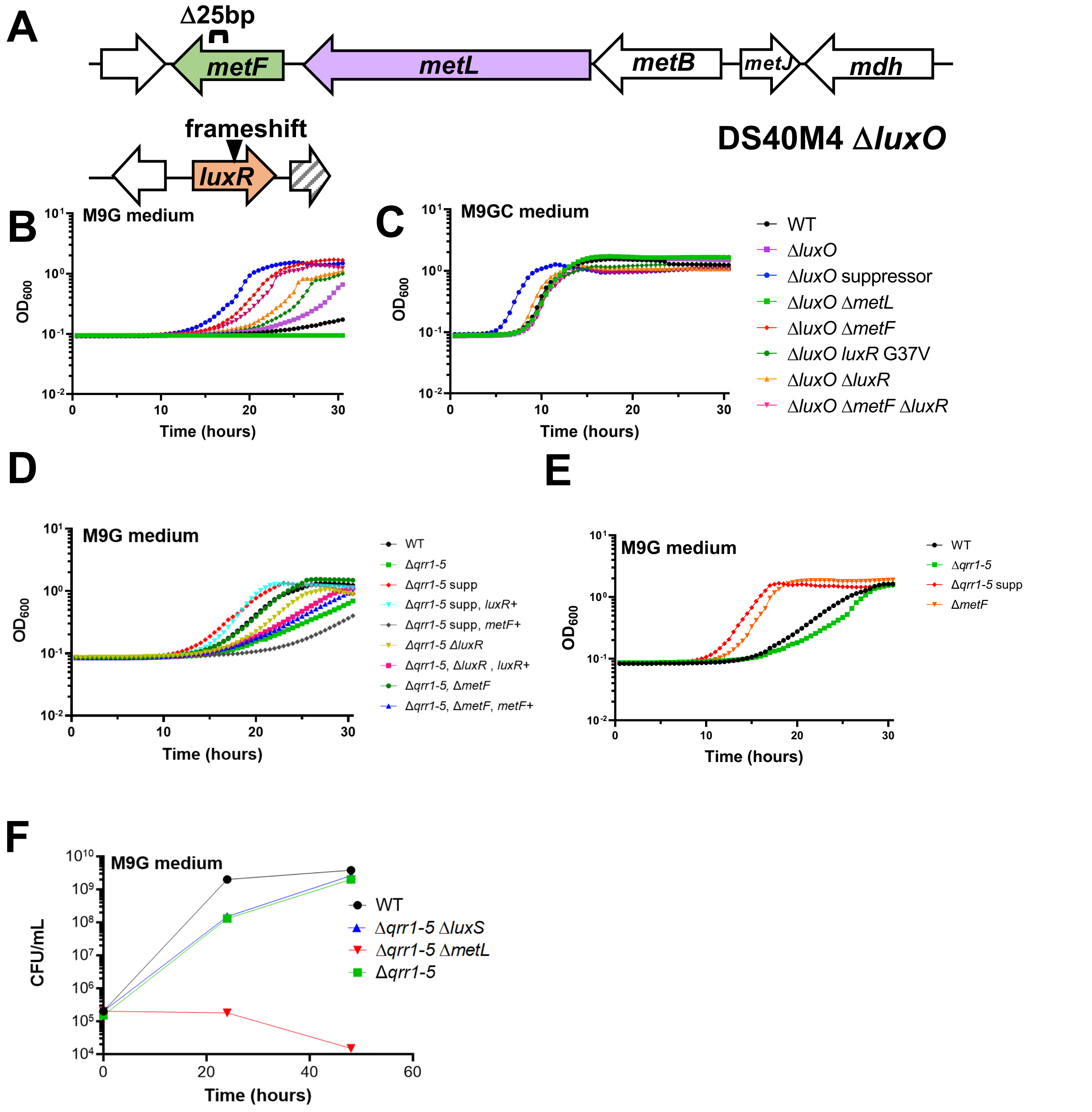

**Figure S2.** (A) Mutations in a Δ*luxO* suppressor mutant strain. (B, C, D, E) Growth curves for *V. campbellii* DS40M4 strains. The y-axis represents cell density $OD_{600}$. (D) The (+) indicates chromosomal complementation of gene at non-native locus. (F) Viable cell counts of DS40M4 strains washed and diluted into M9G medium. For panels B-F, the data shown are from a single experiment that is representative of at least three independent biological experiments for each strain.

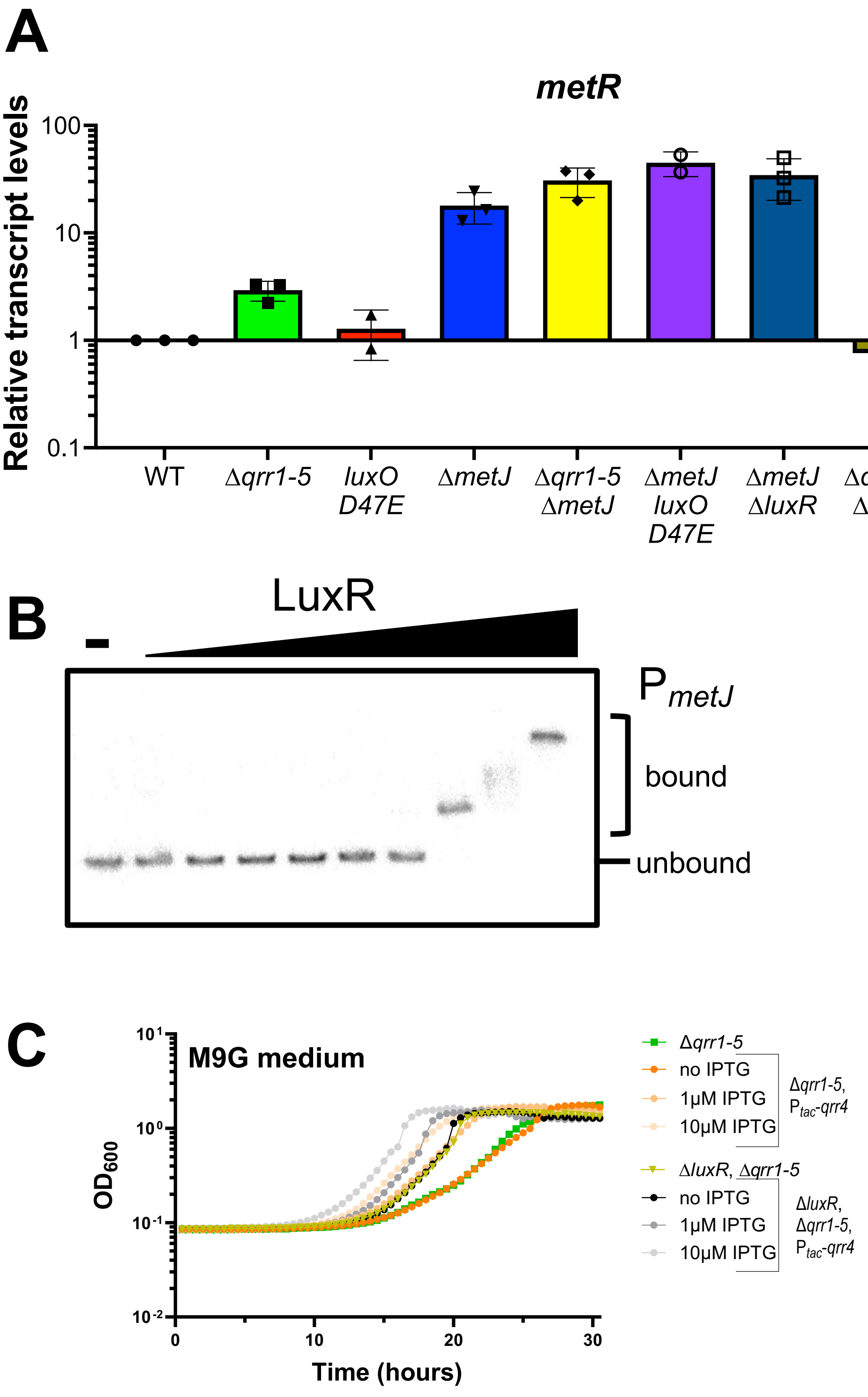

**Figure S3.** (A) Relative *metR* transcript levels determined by RT-qPCR for cultures grown in M9GC. The data could not be determined to be normally distributed (*n* too small), and no significant differences were determined using Dunn's multiple comparisons non-parametric test. Error bars show the mean and standard deviation of three biological replicates. (B) EMSAs with purified WT LuxR with radiolabeled DNA substrate P*metJ* region (ZC011 and ZC012). Protein concentrations are 0.000005, 0.00005, 0.0005, 0.005, 0.05, .5, 5, 50, and 500 nM, compared to no protein control ("-"). This gel is representative of three assays performed with three individual protein preps. (C) Growth curves for *V. campbellii* DS40M4 strains in M9G medium. The y-axis represents cell density $OD_{600}$. The data shown are from a single experiment that is representative of at least three independent biological experiments for each strain under every condition.

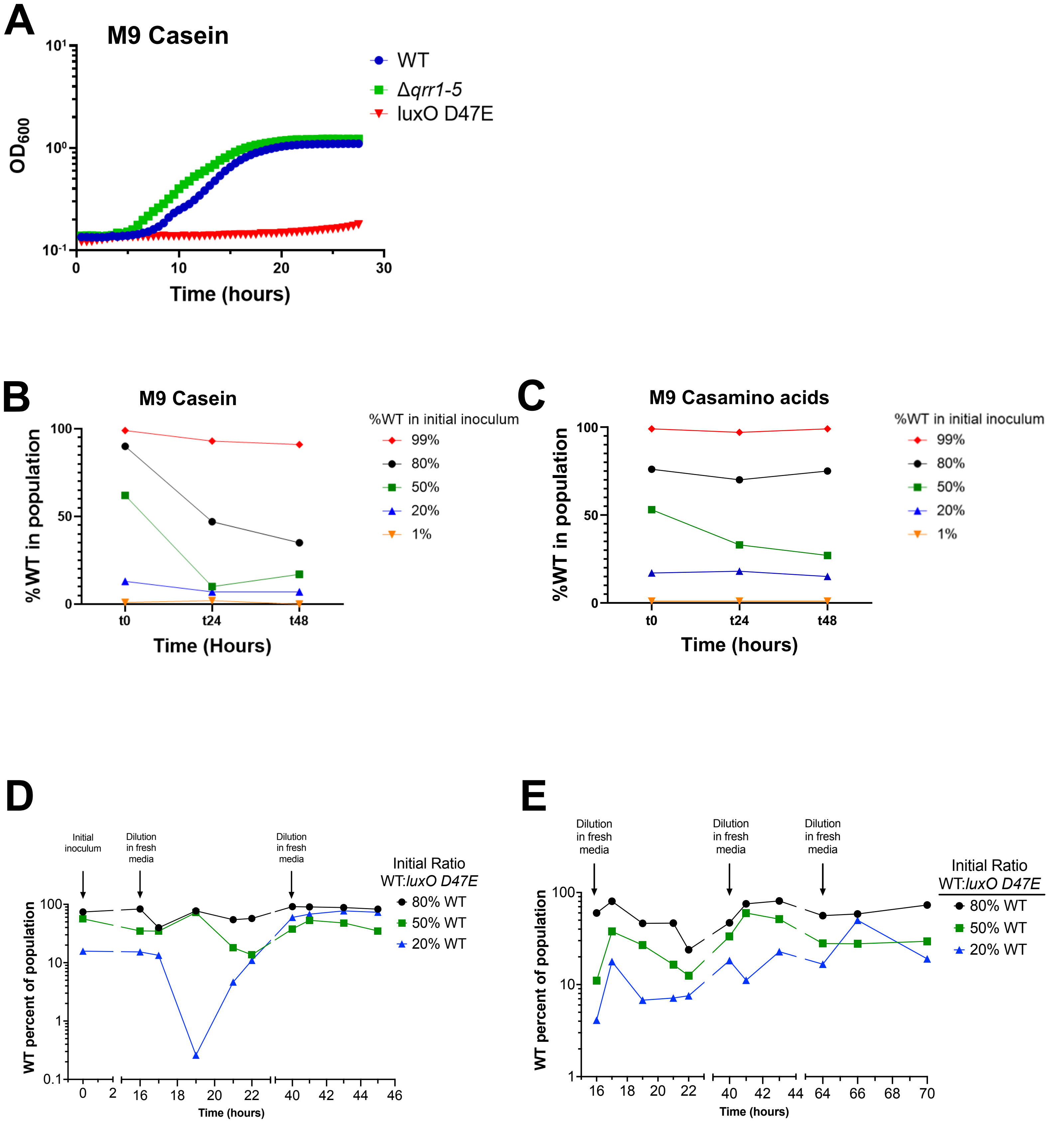

**Figure S4.** (A) Growth assay in M9 medium + 0.5% casein (M9Casein). The y-axis represents cell density $OD_{600.}$ (B, C) Wild-type (marked with Tm) and *luxO D47E* (marked with Spec) strains of DS40M4 were inoculated at initial frequencies of 1-99% wild-type (WT) in M9Casein medium(B) or M9Casaminoacids medium (C). The data in A-C are from a single experiment that is representative of at least three independent biological experiments (D, E) Replicates of competition experiments with wild-type and *luxO D47E* strains. Wild-type (marked with Tm) and *luxO D47E* (marked with Spec) strains of DS40M4 were inoculated at initial frequencies of 80%, 50%, or 20% wild-type (WT) in M9G medium and grown to near stationary phase at t=16 hours, and then diluted to maintain growth in log-phase. Colony forming units (cfus) were measured at each timepoint using selective antibiotics for each strain. The total population number was calculated, and the data are represented as the percent of WT cells in the total population of WT and *luxO D47E*.

**Table S1.** Strains used in this study.

| Strains | Genotype | Reference |
|---|---|---|
| ***V. campbellii* strains** | | |
| BB120 | BB120, wild-type (ATCC BAA-1116) | (1) |
| JAF78 | BB120, Δ*luxO::CM*$^{R}$ | (2) |
| JAF548 | BB120, *luxO* D47E*::KanR* | (3) |
| KT282 | BB120, Δ*qrr1-5* | (4) |
| cas107 | DS40M4, Δ*luxB::SpecR* | (5) |
| cas251 | DS40M4, Δ*qrr1-5* | This study |
| cas197 | DS40M4, Δ*luxO* | This study |
| BP60 | DS40M4, *luxO* D47E | (5) |
| cas537 | DS40M4, Δ*qrr1-5* suppressor mutant | This study |
| cas547 | DS40M4, Δ*qrr1-5*, ΔmetL | This study |
| cas548 | DS40M4, Δ*qrr1-5*, ΔmetF | This study |
| cas549 | DS40M4, Δ*qrr1-5*, *luxR* G37V | This study |
| cas597 | DS40M4, Δ*qrr1-5*, ΔmetF, *luxR* G37V | This study |
| cas601 | DS40M4, Δ*qrr1-5*, ΔmetF, Δ*luxR* | This study |
| cas632 | DS40M4, Δ*qrr1-5*, Δ*luxR* | This study |
| cas600 | DS40M4, Δ*luxR*, Δ*metF* | This study |
| cas196 | DS40M4, Δ*luxR* | This study |
| cas279 | DS40M4, Δ*luxS* | This study |
| cas554 | DS40M4, Δ*qrr1-5*, Δ*luxS* | This study |
| cas628 | DS40M4, Δ*qrr1-5*, pMMB-P*tac-luxS* | This study |
| cas602 | DS40M4, Δ*metJ* | This study |
| cas603 | DS40M4, Δ*metJ*, *luxO* D47E | This study |
| cas604 | DS40M4, Δ*metJ*, Δ*qrr1-5* | This study |
| cas622 | DS40M4, Δ*metJ*, Δ*luxR* | This study |
| cas173 | DS40M4, Δ*luxB::TmR* | This study |
| cas625 | DS40M4, Δ*luxO* suppressor mutant | This study |
| cas595 | DS40M4, Δ*luxO*, Δ*metL* | This study |
| cas596 | DS40M4, Δ*luxO*, Δ*metF* | This study |
| cas620 | DS40M4, Δ*luxO*, *luxR* G37V | This study |
| cas630 | DS40M4, Δ*luxO*, Δ*luxR* | This study |
| cas631 | DS40M4, Δ*luxO*, Δ*luxR,* Δ*metF* | This study |
| cas662 | DS40M4, Δ*qrr1-5* suppressor mutant, Δ*luxB::TmR*+*metF* compliment | This study |
| cas663 | DS40M4, Δ*qrr1-5* suppressor mutant, Δ*luxB::TmR*+*luxR* compliment | This study |
| cas660 | DS40M4, Δ*qrr1-5*, Δ*MetF*, Δ*luxB::SpecR*+*MetF* compliment | This study |
| cas661 | DS40M4, Δ*qrr1-5*, Δ*luxR*, Δ*luxB::SpecR*+*LuxR* compliment | This study |
| cas419 | DS40M4, Δ*qrr1-5,* Δ*luxB::Ptac-qrr4 TmR* | This study |
| cas666 | DS40M4, Δ*qrr1-5,* Δ*luxR,* Δ*luxB::Ptac-qrr4 TmR* | This study |
| ***E. coli* strains** | | |
| S17-1λpir | Wild-type, mating strain | (6) |
| pZRC009 | BL21(DE3), pZC005 | This study |
| pZRC010 | BL21(DE3), pZC006 | This study |

**Table S2.** Plasmids used in this study.

| Strains | Genotype | Reference |
|---|---|---|
| pCS38 | pMMB67EH, PluxC-luxCDABE | (5) |
| pCS48 | pMMB67EH, P*tac-luxS* | (5) |
| pZRC009 | BL21(DE3), pET28b-LuxR (DS40M4) | This study(Twist Bioscience) |
| pZRC010 | BL21(DE3), pET28b-LuxR G37V (DS40M4) | This study(Twist Bioscience) |

**Table S3.** Oligonucleotides used in this study.

| Name | Sequence | Notes |
|---|---|---|
| **qRT-PCR Primers** | | |
| CAS0682 | gcacatcgatgccttaaaccag | qRT-PCR DS40M4 *metF* F |
| CAS0683 | cgtttcttccatcttctcactgc | qRT-PCR DS40M4 *metF* R |
| CAS0684 | cacttacgtaccctcacctcgt | qRT-PCR DS40M4 *metR* F |
| CAS0685 | ctttgatttggtgtgaaagggcg | qRT-PCR DS40M4 *metR* R |
| CAS0678 | gctgactggaatggcgaatac | qRT-PCR DS40M4 *metJ* F |
| CAS0679 | ttaaaacttttagcgggatagagacgg | qRT-PCR DS40M4 *metJ* R |
| **EMSA Primers** | | |
| JCV369 | tgatgttatttatatttatatcatttaaataa | F $P_{luxC}$ Site H |
| JCV620 | ttatttaaatgatataaatataaataacatca | R $P_{luxC}$ Site H |
| ZC011 | atgagtgtatcctttcacggg | F PCR of 239 bp $P_{metJ}$ |
| ZC012 | tcagccattgcgcacc | R PCR of 239 bp $P_{metJ}$ |
